# Evaluation of CB_2_R expression and pyridine-based radiotracers in brains from a mouse model of Alzheimer’s disease

**DOI:** 10.1101/2022.07.26.501500

**Authors:** Vasil Kecheliev, Francesco Spinelli, Adrienne Herde, Ahmed Haider, Linjing Mu, Jan Klohs, Simon M. Ametamey, Ruiqing Ni

## Abstract

Neuroinflammation plays an important role in the pathophysiology of Alzheimer’s disease. The cannabinoid type 2 receptor (CB_2_R) is an emerging target for neuroinflammation and therapeutics of Alzheimer’s disease. Here, we aimed to assess the alterations in brain CB_2_R levels and evaluate novel CB_2_R imaging tracers in the arcAβ mouse model of Alzheimer’s disease amyloidosis. Immunohistochemical staining for Aβ deposits (6E10), microgliosis (anti-Iba1 and anti-CD68 antibodies), astrocytes (GFAP) and the anti-CB_2_R antibody was performed on brain slices from arcAβ mice 17 months of age. Autoradiography using the CB_2_R imaging probes [^18^F]RoSMA-18-d6, [^11^C]RSR-056 and [^11^C]RS-028 and mRNA analysis were performed in brain tissue from arcAβ and nontransgenic littermate (NTL) mice at 6, 17, and 24 months of age. Specific increased CB_2_R immunofluorescence intensities on the increased number of GFAP-positive astrocytes and Iba1-positive microglia were detected in the hippocampus and cortex of 17-month-old arcAβ mice compared to NTL mice. CB_2_R immunofluorescence was higher in the glial cells inside 6E10-positive amyloid-β deposits than peri-plaque with a low background. *Ex vivo* autoradiography showed that the binding of [^18^F]RoSMA-18-d6 and [^11^C]RSR-056 was comparable in arcAβ and NTL mice at 6, 17 and 24 months. The level of *Cnr2* mRNA expression in the brain was not significantly different between arcAβ and NTL mice at 6, 17 or 24 months. In conclusion, we demonstrated pronounced specific increases in microglial and astroglial CB_2_R expression levels in a model of AD-related cerebral amyloidosis/AD mouse model, emphasizing CB_2_R as a suitable target for imaging neuroinflammation.

## Introduction

Abnormal accumulation of amyloid-beta (Aβ) aggregates in Alzheimer’s disease (AD) leads to a cascade of pathophysiological changes, including neuroinflammation, microvascular alterations, synaptic dysfunction, and neuronal loss. Increased numbers of astrocytes and microglia were observed in the vicinity of Aβ plaques in postmortem AD mouse model brains and patients with AD [18]. Microglia are resident macrophages in the central nervous system (CNS) that are important for maintaining brain homeostasis [18] but have also been implicated in the pathophysiology of AD [8,22]. Recent single-cell sequencing transcriptomics for disease-associated microglia (DAM) represents transcriptionally distinct and neurodegeneration-specific microglial profiles with potential significance in AD signatures, including TREM2, CD33, and ApoE [8,21,51,13].

Positron emission tomography (PET) ligands for translocator protein (TSPO) are the most widely used for detecting neuroinflammation and have shown microglial activation preceding Aβ deposition in several animal models, such as APP23, J20, APPSL70, App^NL-G-F^ and PS2APP mice [45,5]. However, limitations in the complex cellular locations, polymorphisms, and nonspecific binding of TSPO and whether TSPO measures microglial proliferation or activation remain to be addressed [63,26]. Novel specific PET tracers for visualizing microgliosis, especially the disease-associated microglia (DAM) subtype, are highly desired.

In the CNS, cannabinoid type 2 receptors (CB_2_Rs) are mainly expressed on microglia at low levels under physiological conditions and are upregulated in acute inflammatory conditions [7]. CB_2_Rs are essential to induce Toll-like receptor-mediated microglial activation [44]. Activation of CB_2_R offers neuroprotective effects, such as reducing Aβ-induced neuronal toxicity [47,57,25,36,62], suppressing microglial activation [43,10], restoring cognitive capacity [58], and ameliorating novel object recognition in animal models of amyloidosis [27], and is thus of therapeutic interest [25]. The expression levels of CB_2_R in animal models of AD amyloidosis have not been extensively characterized. CB_2_R has been shown to be increased and involved in Aβ pathology in 5×FAD [61,31] and J20 mouse models of AD amyloidosis [24] but reduced in the brains of 3×Tg mice (with both amyloid and tau pathology) and aging C57B6 mice [56]. Several CB_2_R ligands have been developed and evaluated [39], including [^11^C]NE40 [54], [^11^C]A-836339 (MDTC) [9,42], [^18^F]MA3 [4], [^18^F]FC0324 [6], [^18^F]JHU94620 [35], [^18^F]LU13 [14], [^18^F]DM102 [34], [^18^F]CRA13 [17], [^11^C]RS-016 [32], [^11^C]RS-028 [16], [^11^C]RSR-056 [50] and [^18^F]RoSMA-18-d6 [15]. Thus far, only one in-human *in vivo* CB_2_R PET using [^11^C]NE40 [1] in patients with AD and healthy controls has been reported, showing no group difference. Only the tracer [^11^C]A-836339 has been evaluated in an AD animal model: Increased [^11^C]A-836339 uptake was observed in the cortex, cerebellum and whole brain of J20 mice compared to wild-type mice [46]; another [^11^C]A-836339 microPET study showed that the uptake was blockable in the cortex of APP/PS1 mice [19].

The aim of the current study was to assess the alterations in CB_2_R and distribution in the brain of the arcAβ mouse model of AD amyloidosis and to evaluate the recently developed pyridine-derived CB_2_R tracers [^11^C]RS-028, [^18^F]RoSMA-18-d6 and [^11^C]RSR-056, which exhibit subnanomolar affinity and high selectivity towards CB_2_R [40].

## Materials and Methods

### Animals

Twenty transgenic arcAβ mice overexpressing the human APP695 transgene containing the Swedish (K670N/M671L) and Arctic (E693G) mutations under control of the prion protein promoter at 6, 17, and 24 months of age and 20 age-matched nontransgenic littermates (NTLs) of both sexes were used in this study [33,23,37]. The arcAβ mouse model exhibits parenchymal plaque as well as cerebral amyloid angiopathy and shows impaired cerebrovascular functions [38,41]. Paper tissue and red mouse house (Tecniplast®) shelters were placed in cages for environmental enrichment. All experiments were performed in accordance with the Swiss Federal Act on Animal Protection and were approved by the Cantonal Veterinary Office Zurich ZH082/18.

For mRNA and autoradiography, mice were anaesthetized under 5% isoflurane and decapitated. Half of the brain hemispheres from arcAβ mice and NTLs were collected, immediately frozen in liquid nitrogen and stored at -80 °C as described earlier [40]. The other half of the brain hemisphere was embedded in TissueTek, frozen and stored at -80 °C for autoradiography. For immunofluorescence staining, mice were perfused under ketamine/xylazine/acepromazine maleate anesthesia (75/10/2 mg/kg body weight, i.p. bolus injection) with ice-cold 0.1 M phosphate-buffered saline (PBS, pH 7.4) and 4% paraformaldehyde in 0.1 M PBS (pH 7.4), fixed for 2 h in 4% paraformaldehyde (pH 7.4) and then stored in 0.1 M PBS (pH 7.4) at 4 °C.

### mRNA isolation and real-time polymerase chain reaction

Total mRNA isolation was performed according to the protocols of the Isol-RNA Lysis Reagent (5PRIME) and the bead-milling TissueLyser system (Qiagen) [40]. A QuantiTect® Reverse Transcription Kit (Qiagen) was used to generate cDNA. The primers (Microsynth) used for quantitative polymerase chain reaction (qPCR) are summarized in **Suppl. Table 1**. Quantitation of *Cnr2* mRNA expression was performed with the DyNAmo™ Flash SYBR® Green qPCR Kit (Thermo Scientific) using a 7900 HT Fast Real-Time PCR System (Applied Biosystems). The amplification signals were detected in real time, which permitted accurate quantification of the amounts of the initial RNA template over 40 cycles according to the manufacturer’s protocol. All reactions were performed in duplicate within three independent runs, and each reaction was normalized against the expression of beta-actin. Quantitative analysis was performed using SDS Software (v2.4) and a previously described 2^−^ΔΔCt quantification method [29]. The specificity of the PCR products of each run was determined and verified with SDS dissociation curve analysis.

### Immunofluorescence

For immunohistochemical analysis, coronal brain sections (40 μm) were cut around Bregma 0- -2 mm and stained with anti-Aβ antibody 6E10, anti-ionized calcium-binding adapter 1 (Iba1) and anti-CD68 for microgliosis, GFAP for astrocytes and anti-CB_2_R antibody as previously described [20] (**Suppl. Table 2**). Sections were mounted with Prolong Diamond mounting media. The brain sections were imaged at ×20 magnification using an Axio Oberver Z1 slide scanner (Zeiss) using the same acquisition settings for all brain slices and at ×63 magnification using a Leica SP8 confocal microscope (Leica). The images were analysed by a person blinded to the genotype using QuPath and ImageJ (NIH). The colocalization of CB_2_R with plaque (6E10 channel), GFAP+ astrocytes or Iba1+ microglia in the cortex and hippocampus was determined on 63×-magnification images. The amount of CB_2_R immunofluorescence within these masks was determined by measuring the mean CB_2_R intensity as well as its integrated density (defined as the factor of the area and average intensity of said area).

### Radiosynthesis and Autoradiography

[^18^F]RoSMA-18-d6 (affinity Ki = 0.8 nM, CB_2_R/CB_1_R > 12000), [^11^C]RSR-056 and [^11^C]RS-028 were synthesized and purified as described previously [40] and formulated with 5% ethanol in water. The molar activities were 156-194 GBq/μmol, 52.3 GBq/μmol, and 86.7-178 GBq/μmol for [^18^F]RoSMA-18-d6, [^11^C]RSR-056, and [^11^C]RS-028, respectively. The radiochemical purity for all three radioligands was > 99%. Autoradiography was performed as described previously [40]. Dissected mouse brains embedded in TissueTek were cut into 10 μm thick sagittal sections on a cryostat (Cryo-Star HM-560MV; Microm) and stored at -80 °C. For [^18^F]RoSMA-18-d6 and [^11^C]RSR-056 autoradiography, slices were thawed on ice and preconditioned in ice-cold buffer (pH 7.4) containing 50 mM TRIS, 5 mM MgCl_2_, and 0.1% fatty acid-free bovine serum albumin (BSA). The tissue slices were dried and then incubated with 1 mL of the corresponding radioligand (0.5-2 nM) for 15 min at room temperature in a humidified chamber. For blockade conditions, GW405833 (10 μM) was added to the solution containing the radioligand. The slices were washed with ice-old washing buffer (pH 7.4) containing 50 mM TRIS, 5 mM MgCl_2_, fatty acid-free BSA, and ice-old distilled water. For [^11^C]RS-028, an additional 2.5 mM EDTA was added to the incubation and washing buffer. After drying, the slices were exposed to a phosphorimager plate (Fuji) for 30 min, and the film was scanned in a BAS5000 reader (Fuji).

### Statistics

Group comparisons in multiple brain regions were performed by using two-way ANOVA with Sidak’s *post-hoc* analysis (GraphPad Prism 9). Comparisons for CB_2_R inside plaque, peri-plaque and parenchymal were performed by using one-way ANOVA with Tukey’s *post-hoc* analysis. All data are presented as the mean ± standard deviation. Significance was set at \**p* < 0.05.

## Results

### Increased CB_2_R expression with proliferation of microglia and astrocytes in the brains of arcAβ compared to NTL

First, we assessed the regional CB_2_R level, the cellular source and the expression of CB_2_R (mean immunofluorescence on the increase number of glia) in the brains of arcAβ mice and NTL mice at 17 months of age. CB_2_R immunofluorescence intensity was increased approximately 4-10-fold in the cortex (4.45 ± 0.25 vs 0.46 ± 0.25, p < 0.0001), hippocampus (4.82 ± 0.10 vs 0.95 ± 0.27, p < 0.0001), and thalamus (1.87 ± 0.31 vs 0.41 ± 0.05, p < 0.0001) of arcAβ mice compared to NTL mice (**Figs. 1, 2a, b**). The background signal of CB_2_R is low in the parenchyma (outside astrocytes/microglia) (**Figs. 1, 2h**). Colocalization analysis indicated that CB_2_R signal density was upregulated on both Iba1+ microglia (379855.49 ± 35254.48 vs 6486.02 ± 2773.22, p < 0.0001) and GFAP+ astrocytes (250994.60 ± 31974.33 vs 19568.63 ± 12282.96, p < 0.0001) in the brains of arcAβ mice compared to NTL mice (**Fig. 2e**). Further analysis indicated that the CB_2_R mean signal intensity was increased in both Iba1+ microglia (55.25 ± 3.76 vs 182.44 ± 11.10, p < 0.0001) and GFAP+ astrocytes (121.37 ± 11.80 vs 7.02 ± 0.79, p < 0.0001) of arcAβ mice compared to NTL mice (**Fig. 2f**).

**Fig. 1.**
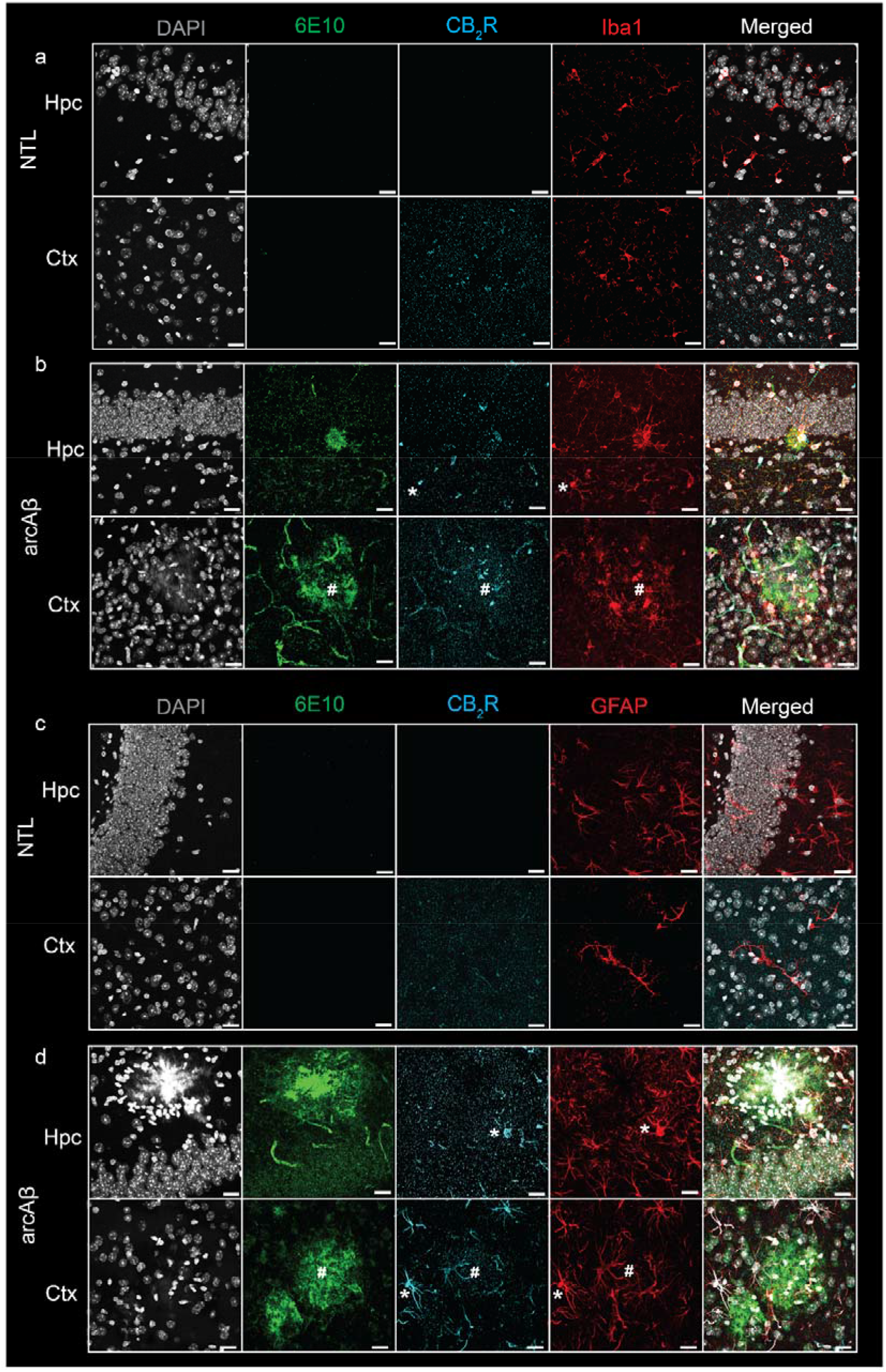
Increased CB_2_R in microglia and astrocytes associated with amyloid-beta deposits in arcAβ mice. **(a-b**) Brain tissue sections of nontransgenic (NTL, n = 3) and arcAβ mice (n = 3) were stained for Aβ (6E10 antibody, green), CB_2_R (cyan), and Iba1 (red) in the hippocampus (Hpc) and cortex (Ctx). Increased CB_2_R and Iba1 immunoreactivity inside and surrounding the plaque. (**c, d**) Staining for Aβ (6E10, green), CB_2_R (cyan), and GFAP (red) in the Hpc and Ctx. Nuclei were counterstained with DAPI (white). Increased CB_2_R and GFAP immunoreactivity inside and surrounding Aβ plaques. * localization of CB_2_R on microglia or astrocytes outside plaque, # colocalization of CB_2_R on microglia or astrocytes within plaque. Scale bar = 20 μm. CB_2_R immunoreactivity was detected on both microglia and astrocytes.

**Fig. 2.**
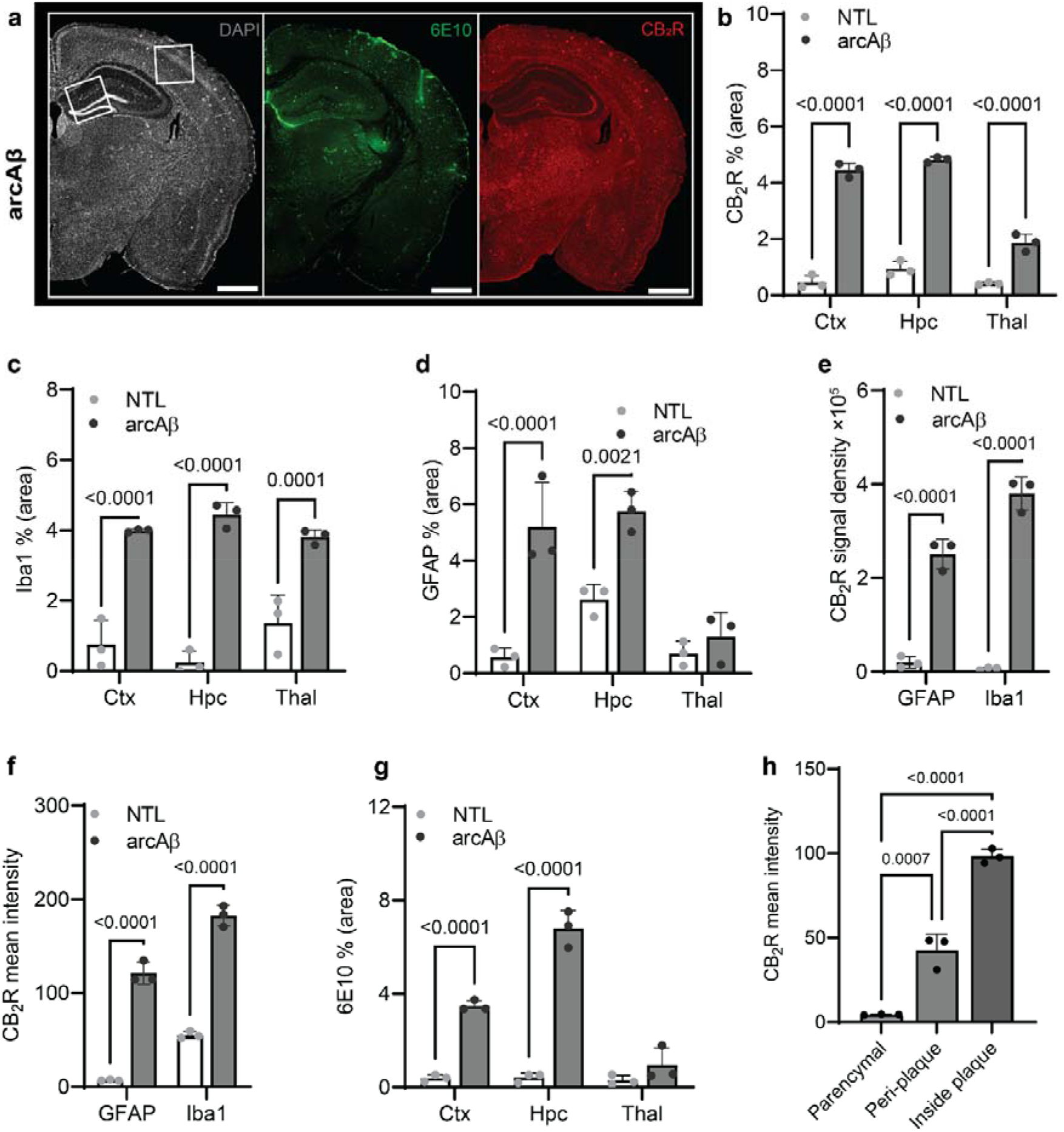
Quantification of microgliosis-, astrocytosis-, CB_2_R- and Aβ plaque-associated enrichment in 17-month-old arcAβ mice. (**a**) Representative CB_2_R (red) and 6E10 (Aβ, green) staining in half hemisphere of one arcAβ mouse brain. (**b**) Increased CB_2_R (% area) in the cortex (Ctx), hippocampus (Hpc), and thalamus (Thal) of arcAβ mice (n = 3) compared to nontransgenic littermates (NTL, n = 3). (**c, d**) Increased levels of Iba1 (% area) in the Ctx, Hpc, and Thal and GFAP (% area) in the Ctx and Hpc of arcAβ mice (n = 3) compared to NTL (n = 3). (**e, f**) Increased CB_2_R signal density and mean signal intensity on both GFAP+ astrocytes and Iba1+ microglia of arcAβ mice (n = 3) compared to NTL (n = 3). (**g**) Increased 6E10 staining of Aβ plaque in the Ctx and Hpc of arcAβ mice (n = 3) compared to NTL (n = 3). (**h**) CB_2_R mean signal intensity on the glia inside plaque is higher than peri-plaque, with low background signal in the parenchymal of arcAβ mice. Scale bar = 20 μm. Data are presented as the mean ± standard deviation.

### Increased CB_2_R associated with 6E10-positive Aβ plaque in the brains of arcAβ compared to NTL

Increased 6E10 immunofluorescence intensity was observed in the cortex (3.48 ± 0.22 vs 0.39 ± 0.15, p < 0.0001) and hippocampus (6.80 ± 0.77 vs 0.42 ± 0.20, p < 0.0001) of arcAβ mice compared to NTL mice and was comparable in the thalamus (0.95 ± 0.73 vs 0.30 ± 0.20, p = 0.2936) (**Figs. 1, 2g**). In the brains of arcAβ mice, CB_2_R immunofluorescence was located on microglia and astrocytes both inside/within plaques (**Fig. 1**). The glial-CB_2_R levels inside the plaques (98.27 ± 4.31 p < 0.0001) and peri-plaques (42.38 ± 9.84, p = 0.0007) were both 20-fold or 10-fold that in the parenchyma (4.5 ± 0.24). The glial-CB_2_R mean fluorescence intensity inside plaque was higher than that located peri-plaque (p < 0.0001) (**Figs. 1, 2h**).

### Increased numbers of Iba1+ and CD68+ microglia and GFAP+ astrocytes in the brains of arcAβ mice compared to NTL mice

Next, we assessed the levels of activated microglia using Iba1 and CD68 and astrocytes using GFAP in the brains of arcAβ mice and NTL mice at 17 months of age. Increased numbers of microglia (Iba1% area) were observed in the vicinity of Aβ plaques and were upregulated in the cortex (3.99 ± 0.04 vs 0.76 ± 0.68, p < 0.0001), hippocampus (4.44 ± 0.35 vs 0.24 ± 0.33, p < 0.0001) and thalamus (3.81 ± 0.21 vs 1.37 ± 0.80, p = 0.0001) of arcAβ mice compared to NTL mice. Increased GFAP % area was associated with plaque in the cortex (5.29 ± 1.57 vs 0.56 ± 0.33, p < 0.0001) and hippocampus (5.75 ± 0.72 vs 2.61 ± 0.53, p = 0.0021) of arcAβ mice compared to NTL mice and was comparable in the thalamus (1.29 ± 0.87 vs 0.70 ± 0.44, p = 0.7933) (**Figs. 1, 2c, d**). CD68 is a lysosomal protein expressed at high levels by activated microglia and at low levels by resting microglia in the CNS. Reactive microglia indicated by increased CD68 surrounding amyloid plaques (6E10) were observed and increased in the cortex (2.74 ± 0.50 vs 1.03 ± 0.29, p = 0.0003) of arcAβ mice compared to NTL mice and were comparable in the hippocampus (1.57 ± 0.40 vs 0.82 ± 0.35, p = 0.0734) and thalamus (1.61 ± 0.19 vs 1.10 ± 0.37, p = 0.2874) (**Fig. 3**).

**Fig. 3.**
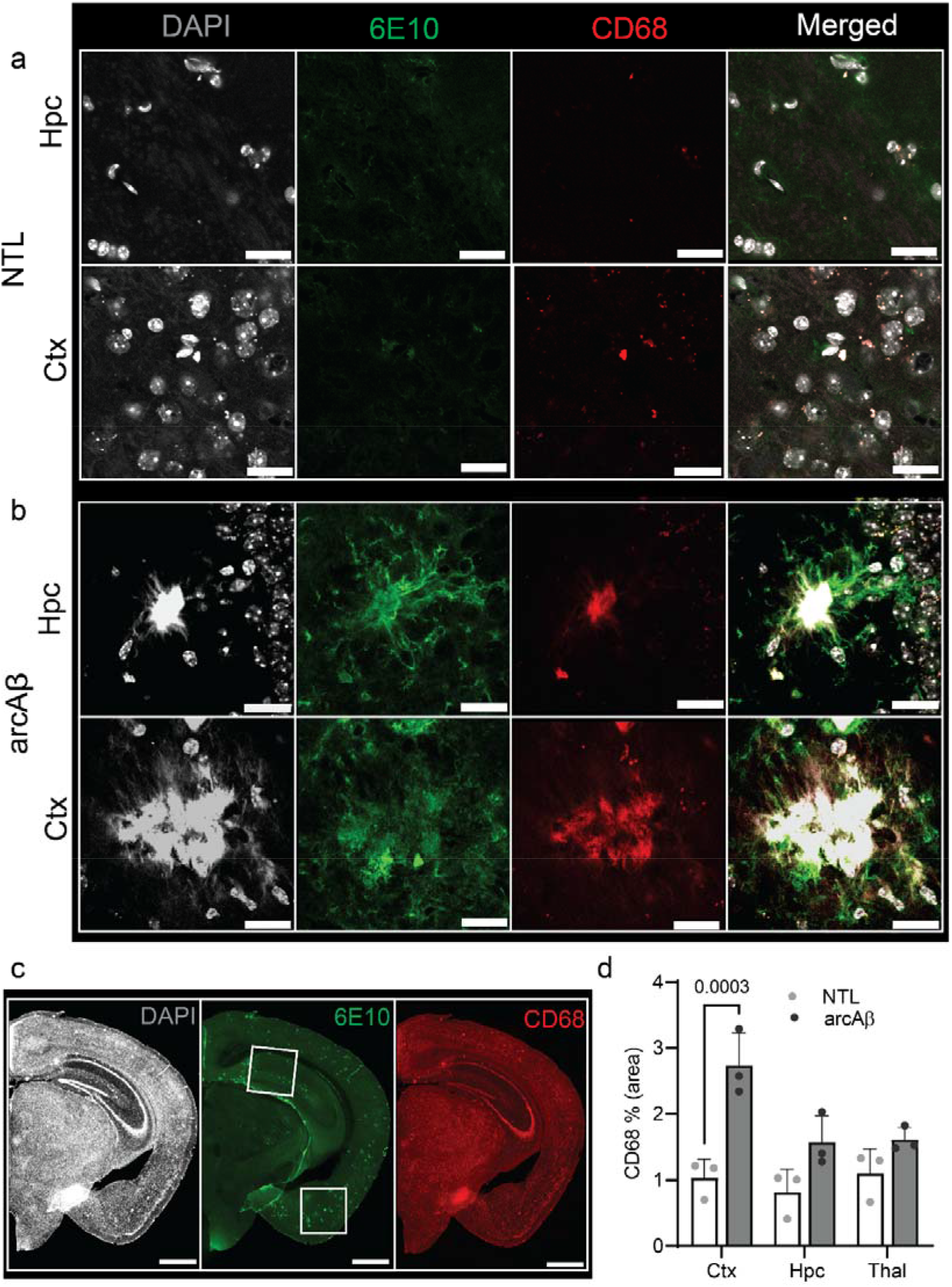
Microgliosis in the arcAβ mouse brain. **(a-c**) Brain tissue sections of nontransgenic (NTL, n = 3) and arcAβ mice (n = 3) were stained for 6E10 (green)/CD68 (red) in the hippocampus (Hpc) and cortex (Ctx). Nuclei were counterstained with DAPI (white). Scale bar = 20 μm. (**d**) Quantification of CD68 signals in the Hpc, Ctx and thalamus (Thal) of arcAβ mice compared to NTL mice confirmed microgliosis in arcAβ mice.

### No difference in whole-brain levels of [^18^F]RoSMA-18-d6 and [^11^C]RSR-056 binding or *Cnr2* expression between arcAβ and NTL mice of different ages

Autoradiography using [^18^F]RoSMA-18-d6, [^11^C]RSR-056 and [^11^C]RS-028 was performed on sagittal brain tissue slides from arcAβ mice to assess the specificity of the probes. The blockage was less than 50% in the brain tissue slices due to the limited number of binding sites, which is much lower than the reported % blockage in the spleen with a high level of CB_2_R binding sites [16,50,15]. [^18^F]RoSMA-18-d6 (40.3 ± 9.2%) showed a higher percentage of specific binding than [^11^C]RSR-056 (32.0 ± 7.8%) and [^11^C]RS-028 (32.0 ± 12.8%, **SFig. 1**).

Thus, we chose [^11^C]RSR-056 and [^18^F]RoSMA-18-d6 to further examine the CB_2_R levels in arcAβ and NTL at 6, 17, and 24 months of age by autoradiography of mouse brain slices. As no specific regional pattern of binding was observed, we analysed the binding level using the whole hemisphere region-of-interest. No difference was observed in brain [^18^F]RoSMA-18-d6 levels between NTL and arcAβ mice at 6 months (0.19 ± 0.03 vs 0.18 ± 0.06 pmol/g tissue, n = 5, 6), 17 months (0.22 ± 0.01 vs 0.19 ± 0.01 pmol/g tissue, n = 3, 5), and 24 months (0.20 ± 0.04 vs 0.22 ± 0.06 pmol/g tissue, n = 5, 5) (**Figs. 4a-b**). Similarly, for [^11^C]RSR-056, no difference in radioactivity accumulation was observed in the brains of NTL and arcAβ mice at 6 months (0.11 ± 0.03 vs 0.12 ± 0.02 pmol/g tissue, n = 5, 6), 17 months (0.18 ± 0.05 vs 0.13 ± 0.02 pmol/g, n = 3, 5), and 24 months (0.14 ± 0.09 vs 0.15 ± 0.03 pmol/g, n = 5, 5) (**Figs. 4c-d**). There was a robust correlation between [^11^C]RSR-056 binding and [^18^F]RoSMA-18-d6 binding in arcAβ and NTL mouse brains (Spearman rank, r = 0.8042, p = 0.0025) (**Fig. 4e**).

**Fig 4.**
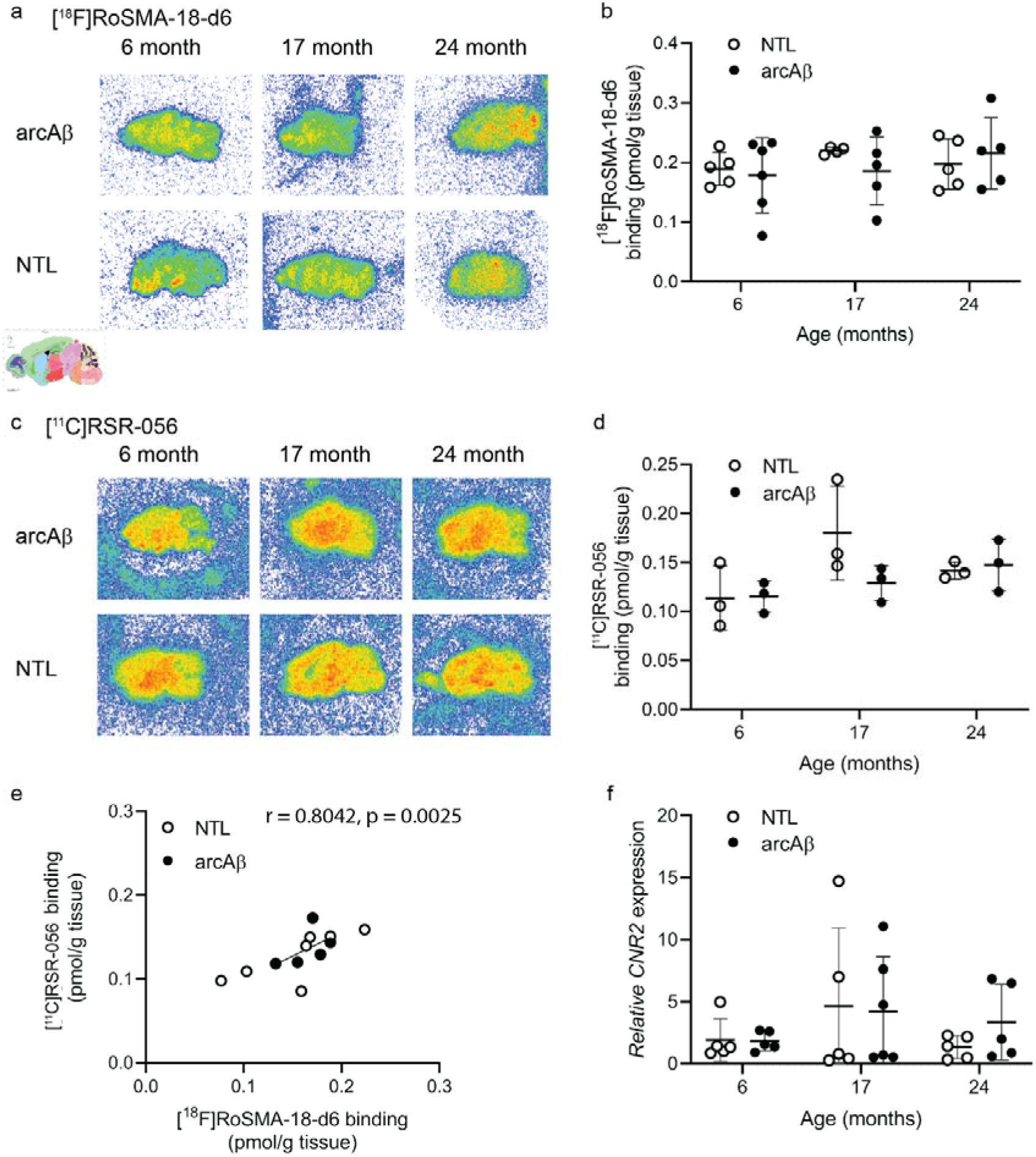
Comparable regional [^18^F]RoSMA-18-d6 and [^11^C]RSR-056 binding in the brains of arcAβ and nontransgenic mice. (**a, c**) Representative [^18^F]RoSMA-18-d6 [^11^C]RSR-056 and autoradiographic images of sagittal brain sections of arcAβ and nontransgenic littermate (NTL) mice at 6, 17, and 24 months of age. Two-way ANOVA, arcAβ v.s. NTL. (b, d) Quantification of ^18^[F]RoSMA-18-d6 and [^11^C]RSR-056 binding in the whole sagittal brain slice. (**e**) Robust correlation between [^11^C]RSR-056 binding and [^18^F]RoSMA-18-d6 binding of arcAβ and NTL mouse brain hemisphere (Spearman rank, r = 0.8042, p = 0.0025). (**f**) *Cnr2* expression in arcAβ and NTL mouse brain hemisphere homogenates at different ages.

Next, we evaluated the mRNA expression levels of *Cnr2* in the left hemisphere from arcAβ and NTL at 6, 17 and 24 months of age that were assessed by autoradiography (n = 5-6/age group). No significant difference was observed in *Cnr2* mRNA expression between the NTL and arcAβ mice at 6 months (1.92 ± 1.72 vs 1.83 ± 0.79), 17 months (4.65 ± 6.30 vs 4.20 ± 4.43), and 24 months (1.33 ± 0.92 vs 3.36 ± 3.07) (**Fig. 4f**).

## Discussion

Here, we demonstrated a local increase in local CB_2_R expression levels in arcAβ mice at 17 months of age compared to NTL mice and evaluated novel PET tracers [^11^C]RSR-056 and [^18^F]RoSMA-18-d6 for detecting brain CB_2_R changes in arcAβ mice. We found increased CB_2_R fluorescence intensities and numbers of microglia and astrocytes inside/surrounding Aβ plaques in arcAβ mice compared to NTL mice. However, we did not observe any significant difference in CB_2_R levels at the whole-brain level measured either by using autoradiography or by mRNA analysis in arcAβ compared to NTL mice at 6, 17, and 24 months.

CB_2_R has been an emerging target for imaging neuroinflammation partly due to its low expression levels under physiological conditions and upregulation under acute inflammatory conditions [52]. We observed that the CB_2_R fluorescence intensity was greatly increased in arcAβ mice compared to NTL mice and was higher inside plaque than peri-plaque and in the parenchyma of arcAβ mice. This observation is different from a previous publication of a significant increase in CB_2_R intensities compared to the core of plaques (radius ≤ 7 μm) [46]. In addition, recent studies have reported astroglial and neuronal expression of CB_2_R in addition to the expression on microglia by using immunostaining and RNAscope techniques [53,12,59,60,46,11,28], although the results are not fully clear. Based on our staining results, we found that CB_2_R expression on both astrocytes and microglia increased significantly in arcAβ mice compared to the negligible level in NTL mice (**Fig 1, 2**). One earlier study using the AD J20 mouse model showed that CB_2_R was highest on neurons in wild-type mice and was enriched in Iba1+ microglia and GFAP+ astrocytes compared to wild-type mice [46]. As immunohistochemical staining was used, concerns regarding the specificity of CB_2_R antibodies have been raised. Specific neuronal subpopulations of CB_2_R have been shown by using fluorescence in situ hybridization and proximity ligand assays in nonhuman primates [49]. However, several studies also reported that CB2-GFP expression is colocalized with Iba1 staining but not with NeuN or GFAP in CB2-GFP BAC transgenic mice [30] and CB2 EGFP^/f/f^ mice [31].

Although Cnr2 expression in AD APP/PS1 has been reported to be upregulated, great variation between animals and a low fold increase lead to insignificance in comparison [55,48,2,3]. Recent gene expression analysis showed that regional Cnr2 expression differs between male/female APP/PS1 mice [55]. Here, we analysed Cnr2 expression using homogenates of half hemispheres of arcAβ and NTL mice with further dissection. We found no difference in Cnr2 expression between arcAβ and NTL mice of different ages.

For preclinical imaging, high variabilities in imaging of brain CB_2_R levels among animal models of neuroinflammation were reported from previous studies. Upregulated levels of brain CB_2_R have been demonstrated in transient middle cerebral artery occlusion ischemic stroke mice using [^18^F]RoSMA-18-d6 [40] and in senescence-accelerated SAMP10 mice using [^11^C]NE40 [59]. Another study by PET using [^11^C]A-836339 in a lipopolysaccharide-injected rat model did not report changes in tracer uptake following neuroinflammation [42]. MicroPET using [^11^C]A-836339 showed increased uptake in the brain areas with Aβ depositions in a J20 mouse model of AD [39]. In the only reported PET study in patients with AD, Ahmad et al. reported lower CB_2_R availability in Aβ-positive AD patients than in healthy controls assessed by PET using [^11^C]NE40 and [^11^C]PIB, respectively [1]. No relationship between [^11^C]NE40 and cerebral Aβ load was observed in this study.

We found that [^11^C]RSR-056 and [^18^F]RoSMA-18-d6 showed 32% and 40% specific binding in the AD mouse brain, respectively, and there was no difference between arcAβ and NTL mice. One of the difficulties is the low CB_2_R expression level in the brain and the low number of binding sites. Using the same tracers, [^11^C]RS-028 [16], [^11^C]RSR-056 [50] and [^18^F]RoSMA-18-d6 [15], lower nonspecific binding has been shown in postmortem spleen and spinal cord tissues from patients with amyotrophic lateral sclerosis than in those from healthy controls. Further development of CB_2_R tracers of even higher affinity to overcome the low number of binding sites (Bmax) is desired. In addition, as species differences exist regarding CB_2_R brain expression, further studies on postmortem brain tissues from patients with AD will provide information on CB_2_R disease relevance.

Limitations in this study: Autoradiography provides information on the probe binding specificity and identifies potential regions of interest with validation from immunohistochemical characterization. Due to a lack of difference from autoradiography, we did not proceed with the *in vivo* measurements.

In conclusion, increases in CB_2_R immunofluorescence intensity on the glia were detected in the brains of arcAβ mice compared to NTL mice and were associated with Aβ deposits. Further improvement of the binding properties of CB_2_R PET tracers will be needed to detect subtle changes in CB_2_R in an AD animal model.

## Acknowledgements

The authors acknowledge Prof. Roger Schibli, department of Chemistry and Applied Biosciences, ETH Zurich; Dr Marie Rouault at the Institute for Biomedical Engineering, ETH Zurich and Mr Daniel Schuppli at the Institute for Regenerative Medicine, University of Zurich for technical assistance. We thank the ALS foundation for partially funding this project.

## Authors’ contributions

RN, JK, SMA, and AMH designed the study; VK performed the staining and microscopy. LN synthesized the radioligands. FS performed the mRNA analysis, RN performed the autoradiography; VK, FS, LN, RN performed data analysis; and VK, RN wrote the initial manuscript. All authors read and approved the final manuscript.

