## Supplementary Figures and Tables for "Evaluation of CB_2_R expression and pyridine-based radiotracers in brains from a mouse model of Alzheimer’s disease"

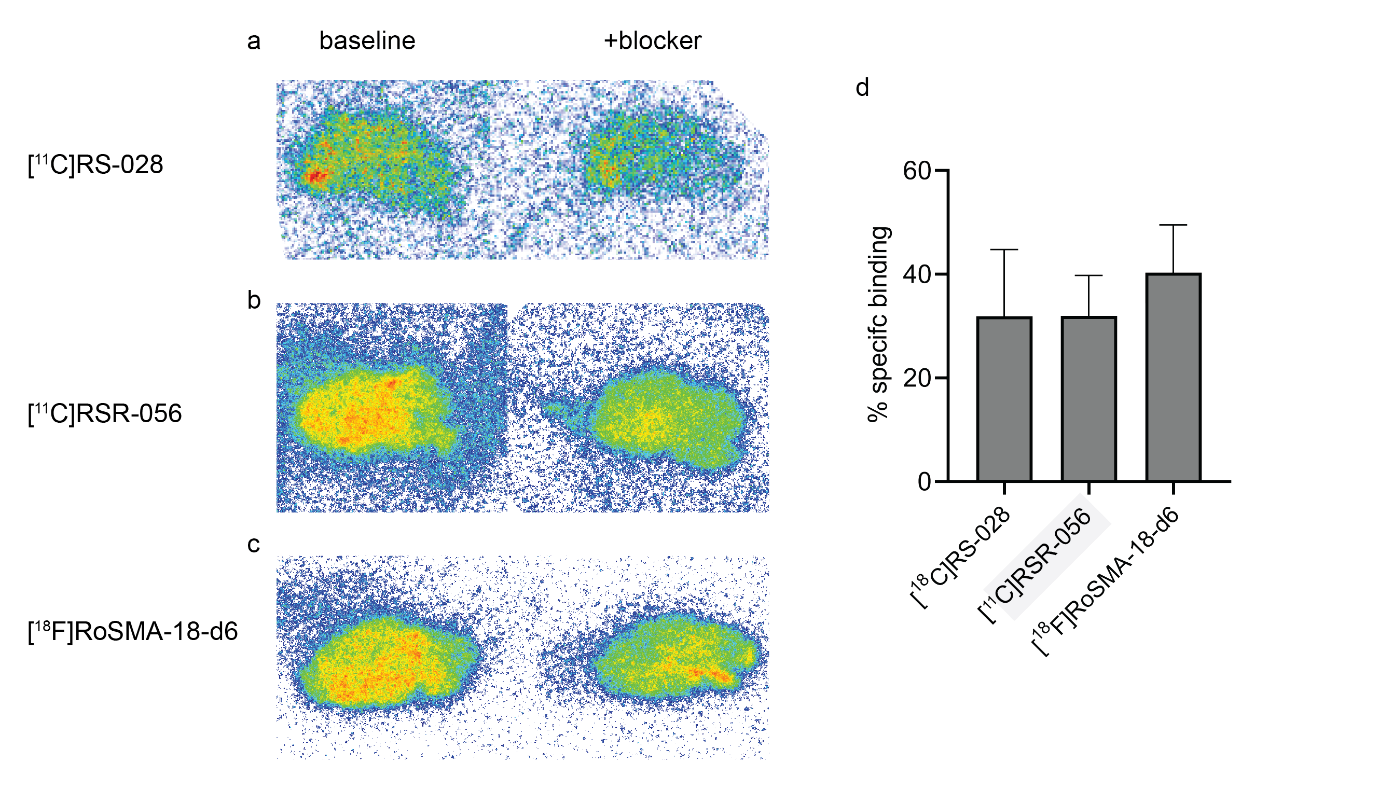

**Supplementary Fig 1. Comparison of different CB_2_R tracers by autoradiography** in mouse brain and in (a-c) Representative [^11^C]RS-028, [^11^C]RSR-056 and [^18^F]RoSMA-18-d6 autoradiographic images of sagittal brain sections of arcAβ mouse and nontransgenic littermates (NTL) mice. (d) Analysis of the percentage of specific binding of [^11^C]RS-028, [^11^C]RSR-056 and [^18^F]RoSMA-18-d6.

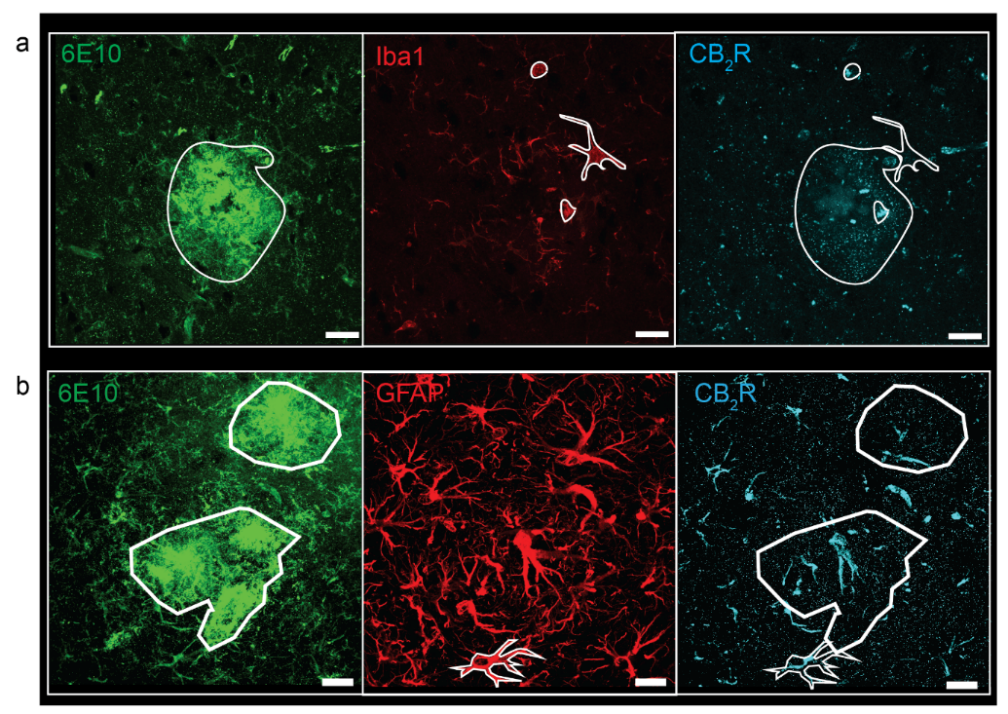

**Supplementary Fig. 2 Illustration of region-of-interest analysis for CB_2_R distribution** (**a, b**) On microglia, astrocytes, inside Aβ plaques of brain tissue sections of arcAβ mice stained using 6E10 (Aβ antibody, green), Iba1 (red) or GFAP (red), and CB_2_R (cyan). Scale bar = 20 μm.

**Suppl. Table 1: Primers used for the quantitative polymerase chain reaction assay on mouse brain tissue**

| mRNA | Primer |
| --- | --- |
| *Actb* | forward 5′-AGACCTCTATGCCAACACAGT-3′, reverse 5′-TGCTAGGAGCCAGAGCAGTAA-3′ |
| *Cnr2* | forward 5′-CTACAAAGCTCTAGTCACCCGT-3′, reverse 5′-CCATGAGCGGCAGGTAAGAAA-3’ |

**Suppl. Table 2 Summary of antibodies and chemicals used in staining**

| **Name** | **Catalog no** | **Dilution** | **Supplier** |
| --- | --- | --- | --- |
| 6E10 | 803001 | 1:1000 | Biolegend |
| CD68 | Ab213363 | 1:1000 | Abcam |
| Iba1 | PA527436 | 1:1000 | Invitrogen |
| Anti-CB_2_R | STJ500369 | 1:1000 | St John’s Laboratory |
| Anti-GFAP | BP5082 | 1:1000 | Acris/Origene |
| Alexa-488 | A32723 | 1:200 | Invitrogen |
| Alexa-555 | A31570 | 1:200 | Invitrogen |
| Alexa-647 | A32795 | 1:200 | Invitrogen |
| DAPI | D1306 | 1:1000 | Sigma–Aldrich |
